# Bisphenol-S and Bisphenol-F alter mouse pancreatic β-cell ion channel expression and activity and insulin release through an estrogen receptor ERβ mediated pathway

**DOI:** 10.1101/2020.09.30.319988

**Authors:** Laura Marroqui, Juan Martinez-Pinna, Manuel Castellano-Muñoz, Reinaldo S. dos Santos, Regla M. Medina-Gali, Sergi Soriano, Ivan Quesada, Jan-Ake Gustafsson, José A. Encinar, Angel Nadal

## Abstract

Bisphenol-S (BPS) and Bisphenol-F (BPF) are current Bisphenol-A (BPA) substitutes. Here we used pancreatic β-cells from wild type (WT) and estrogen receptor β (ERβ) knockout (BERKO) mice to investigate the effects of BPS and BPF on insulin secretion, and the expression and activity of ion channels involved in β-cell function. BPS or BPF rapidly increased insulin release and diminished ATP-sensitive K^+^ (K_ATP_) channel activity. Similarly, 48 h treatment with BPS or BPF enhanced insulin release and decreased the expression of several ion channel subunits in β-cells from WT mice, yet no effects were observed in cells from BERKO mice. PaPE-1, a ligand designed to preferentially trigger extranuclear-initiated ER pathways, mimicked the effects of bisphenols, suggesting the involvement of extranuclear-initiated ERβ pathways. Molecular dynamics simulations indicated differences in ERβ ligand-binding domain dimer stabilization and solvation free energy among different bisphenols and PaPE-1. Our data suggest a mode of action involving ERβ whose activation alters three key cellular events in β-cell, namely ion channel expression and activity, and insulin release. These results may help to improve the hazard identification of bisphenols.

## INTRODUCTION

The relationship between BPA exposure and hormone related diseases (Gore et al., 2015) has raised consumers concern. Consequently, BPA has been progressively substituted by other bisphenol analogs. Among the nearly 15 bisphenol analogs, BPS and BPF are widely consumed and commercialized (Rochester and Bolden, 2015), being the major bisphenol contaminants in indoor dust along with BPA (Liao et al., 2012b). Similar to BPA, the detection frequencies of BPS and BPF were approximately 80% in urine samples collected from the general United States population and several Asian countries (Liao et al., 2012a; Ye et al., 2015). In the United States population, the detection frequency of BPS in urine has increased between 2000 and 2014 while that of BPA trends to decrease since 2010. BPA had a frequency and geometric mean concentrations of 74−99% and 0.36−2.07 μg/L, followed by BPF 42−88%, 0.15−0.54 μg/L and BPS, 19−74%, < 0.1−0.25 μg/L (Ye et al., 2015). BPA has a tolerable daily intake (TDI) determined by the European Food Safety Authority in 2015 of 4 µg/kg-day and in 2017 it was identified by the European Chemical Agency as a substance of very high concern due to its endocrine disrupting properties (Beausoleil et al., 2018). Of note, TDIs for other bisphenols do not yet exist.

BPA has been considered a risk factor in the etiology of type 2 diabetes (T2D) (Alonso-Magdalena et al., 2006; Ropero et al., 2008; Nadal et al., 2009; Batista et al., 2012). Epidemiological and prospective studies associated BPA exposure with alterations in glucose homeostasis or T2D incidence, independently of obesity or other traditional factors (Lang et al., 2008; Shankar and Teppala, 2011; Beydoun et al., 2014; Ranciere et al., 2019). Recent epidemiological data associated BPS urine levels with T2D development in a case-cohort study (Ranciere et al., 2019) and a case-control study (Duan et al., 2018). BPF has been recently associated with abdominal obesity in children (Jacobson et al., 2019), but association with T2D is still unclear.

T2D occurs due to a progressive loss of sufficient β-cell insulin secretion frequently on the background of insulin resistance (American Diabetes, 2018). The use of animal and cellular models indicated a link between BPA exposure and diabetes development (Nadal et al., 2009; Alonso-Magdalena et al., 2011; Le Magueresse-Battistoni et al., 2018). Adult male mice exposed to environmentally relevant doses of BPA presented insulin resistance and hyperinsulinemia in fed state (Alonso-Magdalena et al., 2006; Batista et al., 2012). Furthermore, BPA directly affected β-cell function (Quesada et al., 2002; Alonso-Magdalena et al., 2008; Soriano et al., 2012; Martinez-Pinna et al., 2019). Pancreatic β-cells are excitable cells and, therefore, their electrical activity rules stimulus-secretion coupling. A primary event in the mechanism of insulin release is the blockade of the ATP-sensitive K^+^ (K_ATP_) channels, which control β-cell resting membrane potential. This blockade leads to a typical electrical activity pattern consisting of bursts of action potentials produced by the opening of voltage-gated, Ca^2+^, Na^+^ and K^+^ channels, as well as a rise in intracellular Ca^2+^, which culminates in insulin exocytosis (Rorsman and Ashcroft, 2018). Changes in the expression and/or function of these ion channels result in altered insulin secretion and constitute a serious risk factor for T2D (Hiriart et al., 2014; Jacobson and Shyng, 2020).

Even though BPA may act through different modes of action, it is considered a xenoestrogen able to bind to ERβ and ERα (Wetherill et al., 2007). Both ERs exert their actions through nuclear- and extranuclear-initiated pathways. The nuclear-initiated pathway consists of the direct binding of the ligand bound-ERs to estrogen response elements, which are located in the regulatory regions of ER target genes (Smith and O’Malley, 2004; Heldring et al., 2007). Transcriptional regulation also occurs through tethering of ERs to DNA-bound transcription factors AP-1 and Sp-1 (Ascenzi et al., 2006). Conversely, extranuclear-initiated pathways involve the activation of intracellular signaling cascades that will lead to different effects, including transcriptional regulation (Levin and Hammes, 2016). Although the role of extranuclear-initiated events triggered by environmental estrogens remains poorly understood, rodent models and human studies indicate that this pathway may be important to initiate effects at low doses (Alonso-Magdalena et al., 2008; Vinas and Watson, 2013; Acconcia et al., 2015; Nadal et al., 2018).

In β-cells, nanomolar (1-10nM) concentrations of BPA rapidly (within 10 minutes) block K_ATP_ channels and enhance glucose-stimulated insulin secretion (GSIS) in an ERβ-dependent mechanism (Soriano et al., 2009; Soriano et al., 2012). Longer exposures to BPA (48 hours) regulate gene expression of Ca^2+^, Na^+^ and K^+^ channels, altering electrical activity, Ca^2+^ signaling, and insulin release (Villar-Pazos et al., 2017; Martinez-Pinna et al., 2019). In addition, BPA exposure for 48 h also increases β-cell division in vivo as well as in primary cells. These effects are mimicked by ERβ agonists and abolished in cells from ERβ knockout mice (BERKO), which do not express ERβ in β-cells, suggesting that ERβ activation is necessary for BPA effects in pancreatic β-cells (Boronat-Belda et al., 2020).

Here we studied BPS and BPF effects on insulin release, and ion channel expression and activity in β-cells from wild type (WT) and BERKO mice. Because evidence suggested an important role of ERβ via an extranuclear-initiated pathway, we compared effects elicited by bisphenols with those induced by Pathway Preferential Estrogen-1 (PaPE-1), a compound that binds to ERs and acts preferentially through extranuclear-initiated pathways (Madak-Erdogan et al., 2016). Additionally, we performed molecular docking and dynamic simulations of bisphenols, PaPE-1 and E2 bound to the ERβ ligand-binding domain (LDB) to evaluate the consistencies and variances among these ligands at the molecular level.

## MATERIALS AND METHODS

### Chemical substances

Bisphenol-A was obtained from MP Biomedicals (Cat No 155118; Santa Ana, CA, USA). BPS (Cat No 103039), BPF (Cat No 51453), PaPE-1 (Cat No SML1876), and collagenase (Cat No C9263) were obtained from Sigma-Aldrich (Barcelona, Spain). Bisphenols and PaPE-1 were weekly prepared by dissolution in DMSO (used as vehicle).

### Animals, islet culture and dispersed islet cells

All adult male mice were kept under standard housing conditions (12 h light/dark cycle, food *ad libitum*). BERKO mice were generated as described previously (Krege et al., 1998) and supplied by Jan-Ake Gustafsson’s laboratory. Both WT littermates and BERKO mice were acquired from the same supplier and colony. Mice were sacrificed and islets were isolated as previously described (Nadal and Soria, 1997). For patch-clamp experiments, islets were dispersed into single cells and plated on glass coverslips as described before (Valdeolmillos et al., 1992). Cells were kept at 37 °C in a humidified atmosphere of 95% O_2_ and 5% CO_2_ and used within 48 h of culture. Experimental procedures were performed according to the Spanish Royal Decree 1201/2005 and the European Community Council directive 2010/63/EU. The ethical committee of Miguel Hernandez University reviewed and approved the methods used herein (approvals ID: UMH-IB-AN-01–14 and UMH-IB-AN-02-14).

### Glucose-stimulated insulin secretion (GSIS)

GSIS was performed in islets as previously described (Santin et al., 2016) with slight changes. Briefly, islets were preincubated for 1 h in glucose-free Krebs-Ringer solution. Afterward, islets were sequentially stimulated with 2.8, 8.3, and 16.7 mM glucose for 1 h either in the presence or absence of treatments (as described in Figure 1). Insulin release and insulin content were measured in islet-free supernatants and acid ethanol-extracted islets lysates, respectively, using a mouse insulin ELISA kit (Mercodia, Uppsala, Sweden).

**Figure 1.**
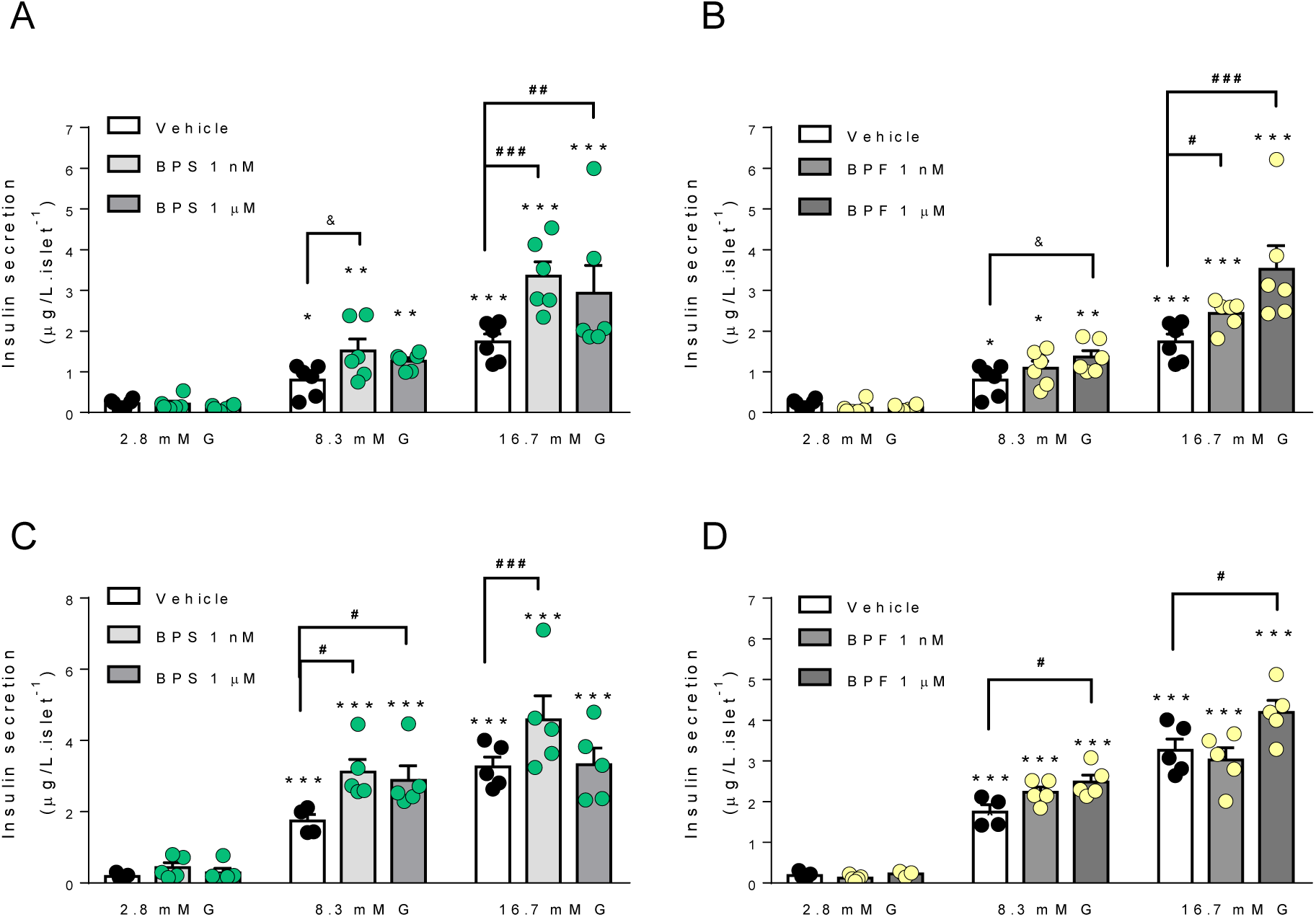
BPS and BPF increase glucose-stimulated insulin secretion in mouse islets. **(A-D)** Insulin secretion was measured at 2.8, 8.3 and 16.7 mM glucose in islets from C57BL/6J mice treated *ex vivo* with vehicle (control; black circles and white bars), 1 nM BPS (green circles and light grey bars) or BPF (yellow circles and light grey bars), or 1 µM BPS (green circles and dark grey bars) or BPF (yellow circles and dark grey bars). **(A and B)** After 2 h of recovery, treatments (vehicle, BPS or BPF) were added to each glucose solution so that the islets remained under treatment during the whole experiment. **(C and D)** Islets were treated *ex vivo* with vehicle BPS or BPF for 48 h, and then, glucose-stimulated insulin secretion was performed in the absence of treatments. Insulin release was measured by ELISA. Data are shown as means ± SEM of six independent islet preparations isolated on three different days: *p≤0.05, **p≤0.01, ***p≤0.001 vs 2.8 mM; ^#^p≤0.05, ^##^p≤0.01, ^###^p≤0.001 comparisons indicated by bars (one-way ANOVA); ^&^p≤0.05 (Student’s t-test).

### Patch-clamp recordings

K_ATP_ channel activity was recorded using standard patch-clamp recording procedures from isolated β-cells as described previously (Valdeolmillos et al., 1992; Vettorazzi et al., 2016). Around 80-90% of the single cells were identified as β-cells. Currents were recorded using an Axopatch 200B patch-clamp amplifier (Axon Instruments Co. CA, USA). Patch pipettes were pulled from borosilicate capillaries (Sutter Instruments Co. CA, USA) using a flaming/brown micropipette puller P-97 (Sutter Instruments Co. CA, USA) with resistance between 3–5 MΩ when filled with the pipette solutions as specified below. Bath solution contained (in mM): 5 KCl, 135 NaCl, 2.5 CaCl2, 10 Hepes and 1.1 MgCl_2_ (pH 7.4) and supplemented with glucose as indicated. The pipette solution contained (in mM): 140 KCl, 1 MgCl_2_, 10 Hepes and 1 EGTA (pH 7.2). The pipette potential was held at 0 mV throughout recording. K_ATP_ channel activity was quantified by digitizing 60 s sections of the current record, filtered at 1 kHz, sampled at 10 kHz by a Digidata 1322A (Axon Instruments Co. CA, USA), and calculating the mean open probability of the channel (*NP*_*o*_) during the sweep. Channel activity was defined as the product of *N*, the number of functional channels, and *P*_*o*_, the open-state probability. *Po* was determined by dividing the total time channels spent in the open state by the total sample time.

For the patch-clamp recordings of voltage-gated Ca^2+^ currents, the whole-cell patch-clamp configuration was used as described previously (Villar-Pazos et al., 2017). Pancreatic β-cells were identified by size (>5 pF) and the corresponding steady-state inactivation properties of the tetrodotoxin (TTX)-sensitive Na^+^ current. Data were obtained using an Axopatch 200B patch-clamp amplifier (Axon Instruments Co. CA, USA). Patch pipettes were pulled from borosilicate capillaries (Sutter Instruments Co. CA, USA) using a flaming/brown micropipette puller P-97 (Sutter Instruments Co. CA, USA) and heat polished at the tip using an MF-830 microforge (Narishige, Japan). The bath solution contained 118 mM NaCl, 20 mM TEA-Cl, 5.6 mM KCl, 2.6 mM CaCl2, 1.2 mM MgCl2, 5 mM HEPES and 5 mM glucose (pH 7.4 with NaOH). The pipette solution consisted of 130 mM CsCl, 1 mM CaCl2, 1 mM MgCl2, 10 mM EGTA, 3 mM MgATP and 10 mM HEPES (pH 7.2 with CsOH). After filling the pipette with the pipette solution, the pipette resistance was 3–5 MΩ. A tight seal (>1 GΩ) was established between the β-cell membrane and the tip of the pipette by gentle suction. The series resistance of the pipette usually increased to 6–15 MΩ after moving to whole-cell. Series resistance compensation was used (up to 70%) for keeping the voltage error below 5 mV during current flow. Voltage-gated Ca^2+^ currents were compensated for capacitive transients and linear leak using a -P/4 protocol. Data were filtered (2 kHz) and digitized (10 kHz) using a Digidata 1322 A (Axon Instruments Co. CA, USA) and stored in a computer for subsequent analysis using commercial software (pClamp9, Axon Instruments Co. CA, USA). Experiments were carried out at 32–34 °C.

### Quantitative real-time PCR

Total RNA was isolated using the RNeasy Micro Kit (Qiagen) and reverse-transcribed using the High Capacity cDNA Reverse Transcription Kit (Applied Biosystems). Quantitative PCR was performed using the CFX96 Real Time System (Bio-Rad, Hercules, CA) as described previously (Villar-Pazos et al., 2017). *Hprt* was used as housekeeping gene. The CFX Manager Version 1.6 (Bio-Rad) was used to analyse the values, which were expressed as relative expression (2^-ΔΔCt^). The primers used herein have been previously described (Martinez-Pinna et al., 2019).

### Molecular docking and dynamics simulations

Crystallographic structure of the rat ERβ LBD in complex with pure antiestrogen ICI 164,384 [rERβ-ΔH12-LBD; UniProt code: Q62986, Protein Data Bank (PDB) code: 1HJ1] was used for molecular docking and long-time dynamic (1 µs) simulation purposes. The missing residues in the 1HJ1 structure (364-377) and the missing side chains (M242, K255, K269, E326, S363, S378, R379, K380 and K435) were reconstructed after generating a homology model at the Swiss-Model server (Biasini et al., 2014; Galiano et al., 2016). Structure of estradiol-bound rat ERβ LBD in complex with LXXLL motif from NCOA5 (rERβ-LBD; UniProt code: Q62986, PDB code: 2J7X) was used for molecular docking and short-time dynamic (100 ns) simulation purposes. The missing residues in the 2J7X structure (239-241 and 369-374) and the missing side chains (V237, M242, K255, E376, R374, K398, and K426) were reconstructed after generating a homology model at Swiss-Model server (Biasini et al., 2014; Galiano et al., 2016) using the 2J7X structure as a template. Molecular docking and dynamics simulations were carried out using YASARA structure v19.9.17 software as previously described (Encinar et al., 2015; Galiano et al., 2016; Ruiz-Torres et al., 2018). The ligand-protein interactions have been detected with the Protein–Ligand Interaction Profiler (FLIP) algorithm (Salentin et al., 2015). Foldx 5.0-calculated (Delgado et al., 2019) was used for frequency distributions of intermolecular protein interaction energy for the subunits of the rERβ-ΔH12-LBD dimer in the presence of different ligands in each LBD cavity (Figure S5).

### Data analysis

The GraphPad Prism 7.0 software (GraphPad Software, La Jolla, CA, USA) was used for statistical analyses. Data are presented as the mean ± SEM. Statistical analyses were performed using Student’s t-test or one-way ANOVA. p values ≤ 0.05 were considered statistically significant.

## RESULTS

### BPS and BPF affect insulin release

Previous data indicate that treatment with 1 nM BPA rapidly enhances insulin secretion in islets from mice and humans (Alonso-Magdalena et al., 2006; Soriano et al., 2012). To investigate whether BPS and BPF would have similar effects, we treated islets during 1 h with two concentrations of BPS and BPF (1 nM and 1 µM), and we measured insulin release in response to different glucose concentrations (2.8, 8.3 and 16.7 mM). Exposure to 1 nM and 1 µM BPS enhanced GSIS at stimulatory glucose concentration, mainly at 16.7 mM (**Figure 1A**). Regarding BPF, we observed a slight increase at 1 nM that was significant only in the presence of 16.7 mM glucose. BPF 1 µM, however, increased GSIS at both 8.3 and 16.7 mM glucose (**Figure 1B**). We used BPA as a positive control and 1 nM BPA increased GSIS, as expected (**Figure S1A**). Of note, insulin content remained unchanged upon treatment with BPS, BPF, and BPA (**Figure S1C-E**). Longer BPA treatment (48 h) induced insulin hypersecretion in response to stimulatory glucose concentrations (Alonso-Magdalena et al., 2008; Villar-Pazos et al., 2017). We then investigated whether treatment with BPS or BPF during 48 h would also change GSIS. BPS at 1 nM and 1 µM enhanced insulin secretion in response to 8.3 mM glucose. However, when glucose concentration was increased to 16.7 mM, BPS was effective at 1 nM but ineffective at 1 µM (**Figure 1C**). When the same experiment in Figure 1C was performed with BPF, we only observed a potentiation of insulin release at 1 µM at stimulatory glucose concentrations (**Figure 1D**), which indicated a more potent action of BPS compared to BPF. Treatment with BPA, BPS or BPF did not modify insulin content (**Figure S1F-H**).

### BPS and BPF diminish K_ATP_ channel activity via ERβ

We have previously demonstrated that acute BPA treatment potentiated GSIS after decreasing K_ATP_ channel activity (Soriano et al., 2012). Moreover, BPA effects, which were not observed in cells from BERKO mice, were reproduced by the endogenous ligand, 17β-estradiol (E2), as well as the ERβ agonist diarylpropionitrile (DPN) (Soriano et al., 2009; Soriano et al., 2012). Acute treatment with BPS induced a rapid increase in heart rate in response to catecholamines (Gao et al., 2015). BPS also rapidly depressed left ventricular contraction and myocyte contractility (Ferguson et al., 2019). In both cases, the ERβ antagonist PHTPP abolished BPS actions, suggesting the involvement of ERβ (Gao et al., 2015; Ferguson et al., 2019).

To assess whether acute exposure to BPS or BPF would modulate K_ATP_ channel activity, we performed patch-clamp recordings in the cell-attached mode in dispersed β-cells from WT and BERKO mice (**Figure 2**). Treatment with 1 nM BPS during 10 minutes was enough to decrease K_ATP_ channel activity by 35% (**Figure 2A,B**), whereas no effects were observed in cells from BERKO mice (**Figure 2A,C**). A similar experiment was performed using 1 nM and 10 nM BPF. While 1 nM BPF did not modify K_ATP_ channel activity in β-cells from WT and BERKO (data not shown), 10 nM BPF decreased K_ATP_ channel activity in cells from WT (**Figure 2D**) but not in cells from BERKO mice (**Figure 2E**). These findings indicate that the rapid GSIS enhancement observed in **Figure 1A** may be a consequence of bisphenol-induced K_ATP_ channels closure.

**Figure 2.**
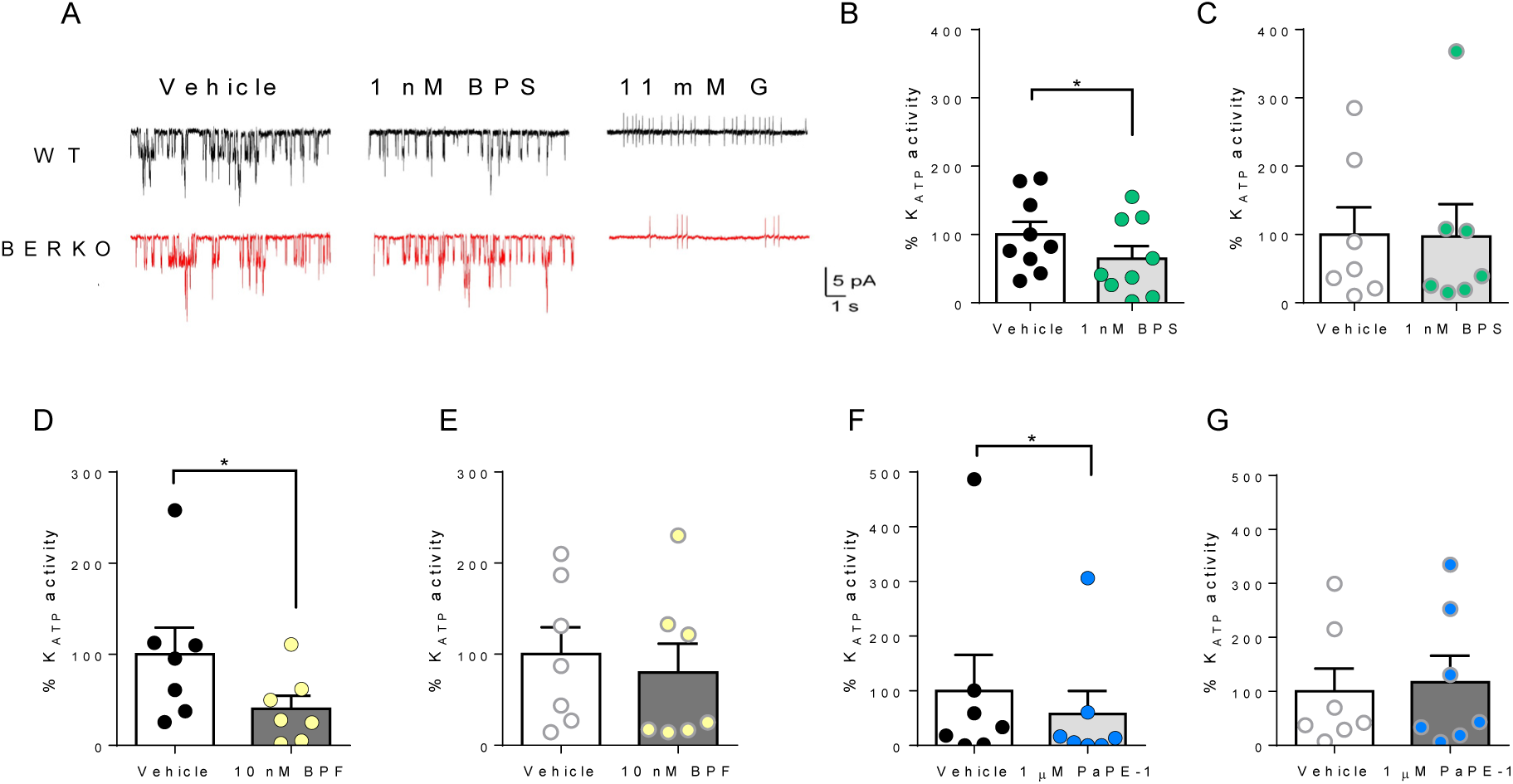
BPS, BPF and PaPE-1 block K_ATP_ channel activity in mouse pancreatic β-cells. **(A)** Representative recordings of K_ATP_ channel activity in β-cells isolated from wild-type (WT) (black traces) or BERKO (red traces) mice in control condition (0 mM glucose; vehicle; left column), in 1 nM BPS (middle column) and in 11 mM glucose (right). Channel openings are represented by downward deflections, reflecting inward currents due to the high K^+^ content of the pipette. The abolition of K_ATP_ activity and generation of action currents at 11 mM glucose was used as a positive control of pancreatic β-cell identity (right column). **(B-G)** Quantification of the K_ATP_ channel activity in β-cells isolated from wild-type (WT) **(B, D and F)** or BERKO **(C, E and G)** mice treated *in vitro* with vehicle (black or white circles and white bars), 1 nM BPS (**B and C**; green circles and light grey bars), 10 nM BPF (**D and E**; yellow circles and dark grey bars) or 1 μM PaPE-1 (**F and G**; blue circles and black bars). The effect of three bisphenols was measured after 7±1 min of acute application. Data are represented as a percentage of activity with respect to resting conditions (0 mM Glucose). Experiments were carried out at 32-34°C. Data are shown as means ± SEM of the number of cells recorded in WT (n=7-9 cells) and BERKO (n=7-9 cells) mice. These cells were isolated from three mice on three different days. *p≤0.05 vs control (Student’s paired t-test).

The fact that this is a rapid action, occurring minutes upon treatment, indicates that low concentrations of bisphenols trigger a non-genomic action via an extranuclear-initiated pathway, likely by binding to ERβ. To test this hypothesis we used PaPE-1, a new ERα and ERβ ligand that acts preferentially through extranuclear-initiated pathways (Madak-Erdogan et al., 2016). Treatment with 1 µM PaPE-1 decreased K_ATP_ channel activity in cells from WT mice (**Figure 2F**) but had no effect in cells from BERKO mice (**Figure 2G**). Of note, 1 nM PaPE-1 did not change K_ATP_ channel activity (data not shown). These results emphasize that PaPE-1 triggers a rapid extranuclear-initiated pathway via ERβ in β-cells.

### Bisphenols downregulate ion channel subunits gene expression

Stimulus-secretion coupling in β-cells depends on the electrical activity generated by ion channels. BPA treatment for 48 h decreased the mRNA expression of genes encoding Ca^2+^ (*Cacna1e*), K^+^ (*Kcnma1* and *Kcnip1*) and Na^+^ (*Scn9a*) channel subunits, which might explain, at least in part, the BPA-induced alteration in GSIS (Villar-Pazos et al., 2017; Martinez-Pinna et al., 2019). In Figure 3 A,E,I) we used BPA as a control to probe that in this preparation it decreased *Cacn1e, Kcnma1, Kcnip* and *Scn9* as already described (Villar-Pazos et al., 2017; Martinez-Pinna et al., 2019). We found that BPS modulated *Cacna1e* mRNA expression in a non-monotonic dose response (NMDR)-dependent manner: exposure to 1 nM BPS reduced *Cacna1e* mRNA expression by 50%, while exposure to 100 nM and 1 µM did not significantly change *Cacna1e* expression (**Figure 3B**). This BPS-induced decrease in *Cacna1e* expression at 1 nM was associated to a reduction in Ca^2+^ currents in cells from WT (**Figure S2A,C,I**), but not in cells from BERKO (**Figure S2B,D,I**) mice. Of note, 100 nM and 1 µM BPS did not modify Ca^2+^ currents in cells from WT or BERKO mice (**Figure S2**). It is very likely that the decrease in Ca^2+^ currents induced by 1 nM BPS is a consequence of *Cacna1e* gene downregulation because both follow the same dose pattern. Regarding BPF, *Cacna1e* mRNA expression was not changed by treatment with 1 nM BPF for 48 h, but it was decreased upon exposure to 100 nM and 1 µM BPF (**Figure 3C**). Measurement of Ca^2+^ currents showed that only exposure to 1 µM BPF significantly decreased Ca^2+^ currents in cells from WT mice (**Figure S3A,G,I**), while no effects were observed in cells from BERKO mice (**Figure S3B,H,J**). Once again, we used PaPE-1 to study the possible involvement of an extranuclear-initiated pathway in the regulation of *Cacna1e* expression. Treatment with 1 µM PaPE-1 decreased *Cacna1e* expression (**Figure 3D**), which indicates that this gene can be regulated by a signaling pathway initiated outside the nucleus.

**Figure 3.**
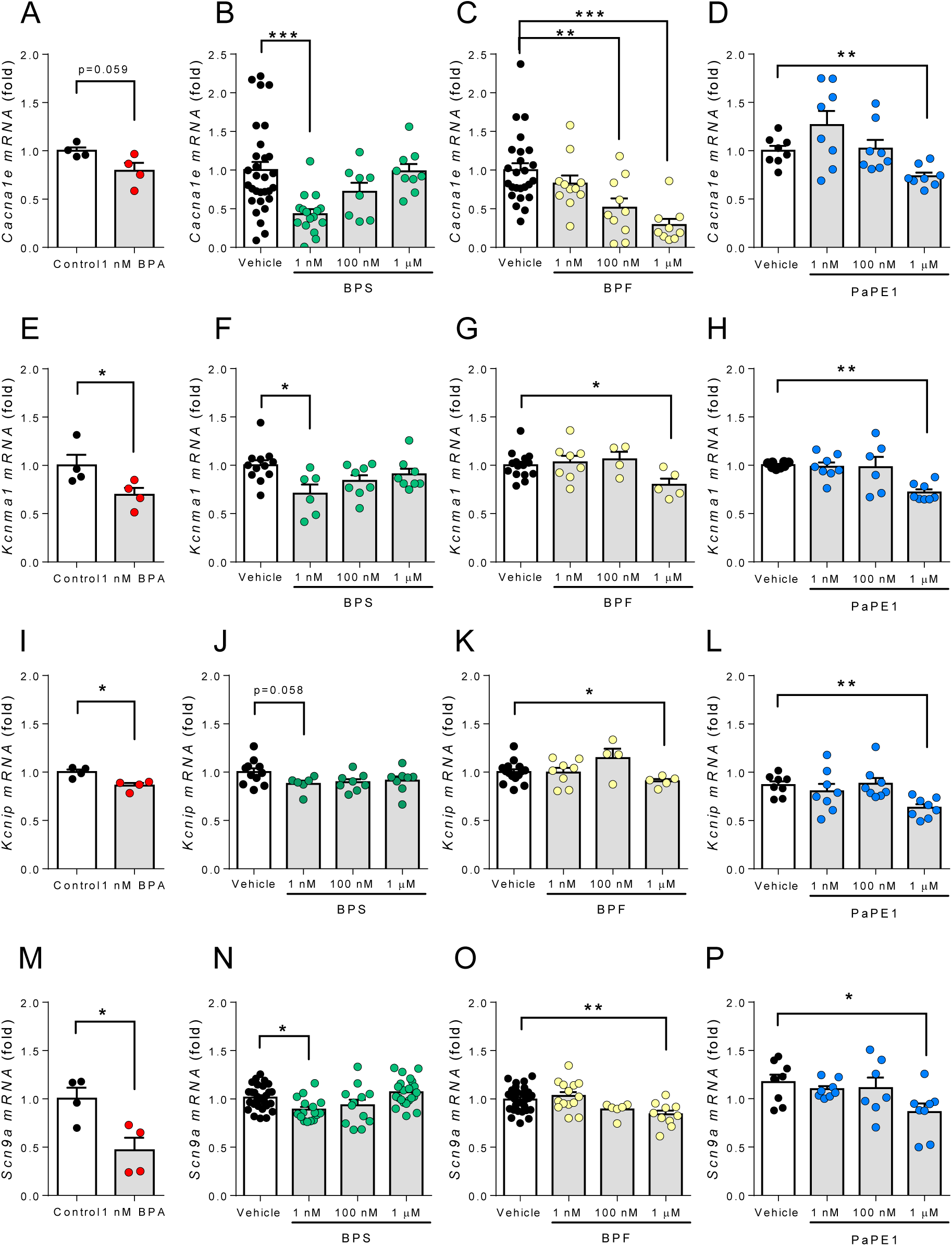
BPA, BPS, BPF and PaPE-1 reduce *Cacna1e, Kcnma1, Kcnip* and *Scn9a* expression in mouse islets. mRNA expression of *Cacna1e* **(A-D)**, *Kcnma1* (), *Kcnip* **(I-L)** and *Scn9a* **(M-P)** was measured in islets from C57BL/6J mice treated *ex vivo* with vehicle (control; black circles and white bars), BPA **(A, E, I and M**; red circles and light grey bars), BPS **(B, F, J and N**; green circles and light grey bars), or BPF **(C, G, K and O**; yellow circles and light grey bars) or PaPE-1 **(D, H, L and P**; blue circles and light grey bars) at 1, 100 and 1000 nM for 48 h. mRNA expression was measured by qRT-PCR and normalized to the housekeeping gene *Hprt1*, and is shown as fold vs. mean of the controls. Data are shown as means ± SEM of four to twenty-nine independent samples from up to twenty-nine islets preparations isolated on at least three different days: *p≤0.05, **p≤0.01, ***p≤0.001 (ANOVA one way).

Like what we observed for *Cacna1e* expression, *Kcnma1* (**Figure 3E-H**), *Kcnip1* (**Figure 3I-L**), and *Scn9a* (**Figure 3M-P**) mRNA expression was downregulated by BPA, BPS, BPF and PaPE-1. BPS decreased *Kcnma1, Kcnip1*, and *Scn9a* at 1 nM in an NMDR manner, while BPF and PaPE-1 were effective at 1 µM (**Figure 3**).

As a negative control, we used 4,4’-(9-fluorenylidene)diphenol, BPFL (also named BHPF), which binds to the androgen receptor and acts as an antiestrogen (Zhang et al., 2017; Keminer et al., 2019). As expected, BPFL treatment at different concentrations did not change ion channel gene expression or Ca^2+^ currents (**Figure S4**).

Overall, these results demonstrate that BPS decreased the transcription of ion channel subunits at concentrations as low as 1 nM, while BPF needed higher concentrations (100 nM and 1 µM) to decrease the expression of the same genes. This effect was mimicked by PaPE-1, suggesting that bisphenols may regulate gene expression via extranuclear ERs.

We previously used β-cells from BERKO mice as well as the ERβ ligand DPN to study the role of ERβ on the regulation of ion channel subunit gene expression induced by BPA (Villar-Pazos et al., 2017; Martinez-Pinna et al., 2019). To evaluate whether ERβ would also play a role in BPS- and BPF-induced regulation of ion channel expression, we incubated islets from WT and BERKO mice with 1 nM BPS or 1 µM BPF for 48 h. Similarly, to the results depicted in **Figure 3**, both 1 nM BPS and 1 µM BPF decreased *Cacna1e, Kcnma1* and *Scn9a* mRNA expression in islets from WT mice (**Figure 4 A-C**). This decrease, however, was abolished in islets from BERKO mice (**Figure 4 D-F**). Notably, 1 µM BPF increased *Cacna1e* expression in BERKO mice, suggesting a role for receptors other than ERβ in the regulation of this gene.

**Figure 4.**
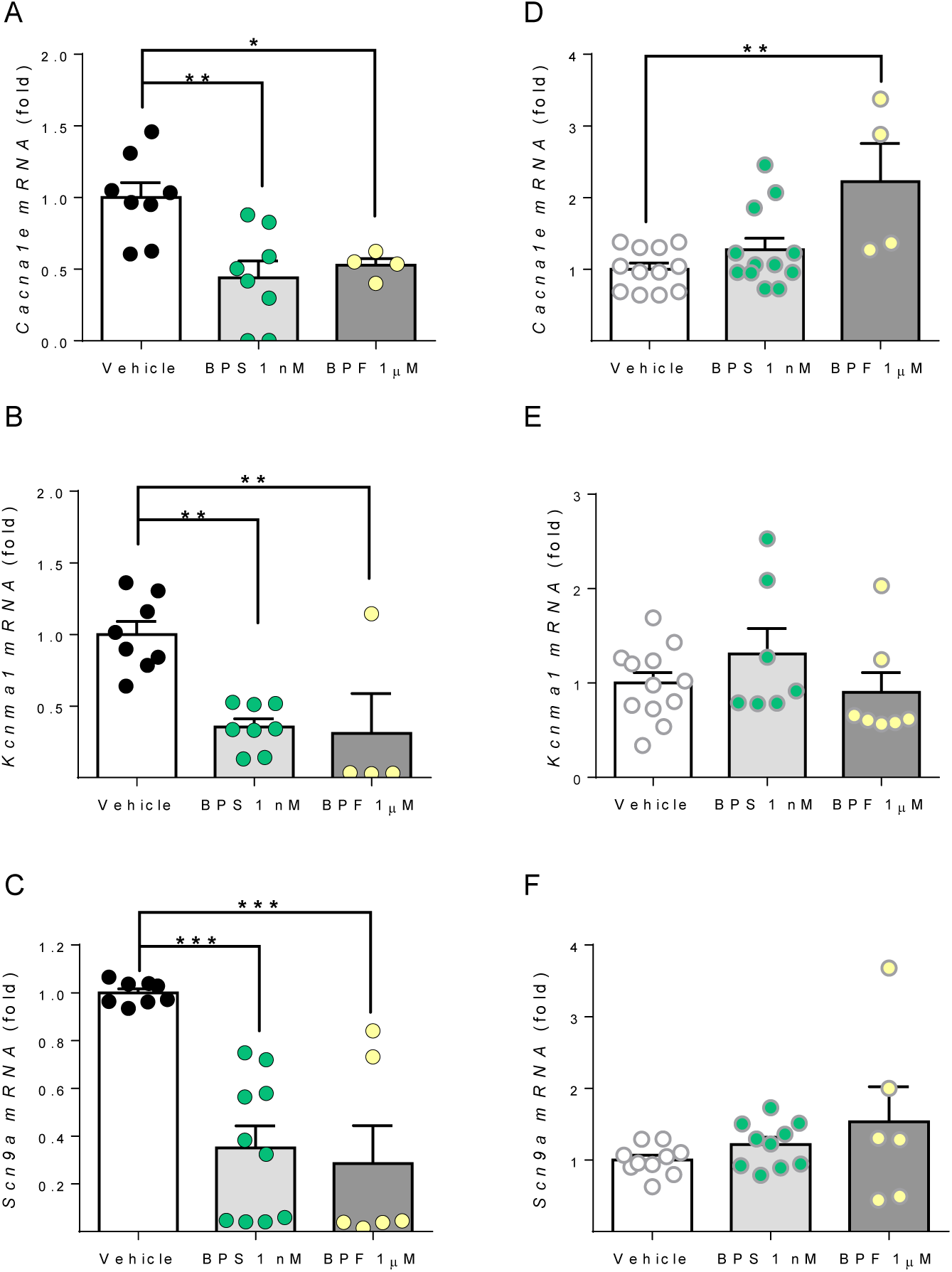
BPS, and BPF, reduce *Cacna1e, Kcnma1* and *Scn9a* expression in islets from wild type but not from BERKO mice. mRNA expression of *Cacna1e* **(A and D)**, *Kcnma1* **(B and E)** and *Scn9a* **(C and F)** in islets isolated from wild-type **(A, B and C)** or BERKO **(D, E and F)** mice treated *ex vivo* with vehicle (control; black circles and white bars), 1 nM BPS (green circles and light grey bars), or 1 µM BPF (yellow circles and dark grey bars) for 48 h. mRNA expression was measured by qRT-PCR and normalized to the housekeeping gene *Hprt1*, and is shown as fold vs mean of the controls. Data are shown as means ± SEM of four to eight independent islet preparations isolated on at least three different days: *p≤0.05, **p≤0.01, ***p≤0.001 (ANOVA one way).

### Molecular dynamics simulations of bisphenols bound to the rat ERβ LBD cavity

To investigate potential modifications in bisphenols and PaPE-1 binding to ERβ ligand binding domin (LBD) that might help to, at least partially, explain the different biological activity of bisphenols observed herein, we performed computational analyses of molecular docking and dynamics simulations.

Despite the numerous studies using human ERα structures on molecular dynamics simulations (Celik et al., 2007; Fratev, 2015; Chen et al., 2016; Jereva et al., 2017; Li et al., 2018; Shtaiwi et al., 2018), the ERβ isoform has not yet been analyzed. Furthermore, ERα LBD and ERβ LBD have not been crystallized in mice, even though the resolved rat structure is well known. Because rat and mouse ERβ LBD sequences differ by only 3 amino acids, including two conservative mutations (**Figure S5A,B**), we chose to use rat structures in our analyses. A recent study using E2, BPA, BPS and BPAF indicated that the root mean square deviation (RMSD) values calculated from the heavy atoms of the ligands might be an important parameter to analyze ligand dynamics particularly implicated in nuclear-initiated events (Li et al., 2018). Therefore, to evaluate if our results fixed with a classic nuclear initiated event, we first studied molecular dynamics simulation of the transactivation helix (H12) closed rERβ-LBD. The natural ligand E2 showed no deviations from the starting configuration for over 100 ns (**Figure S6**), whereas deviations reached 2 Å in the presence of the Src coactivator peptide (**Figure S6**). Meanwhile, rearrangements in conformations are evident from the ligand heavy atom RMSDs in all three bisphenols. In addition, we observed that BPA and BPF showed rapid variations due to faster ring-flipping dynamics (**Figure S6A-C**), similarly to what has been previously shown (Li et al., 2018). The repositioning of H12 in the “mouse trap” conformation decisively influences the MM/PBSA solvation binding energy (Celik et al., 2007). We observed that the solvation binding energy for E2 and BPA was higher than that for BPS and BPF (**Figure S7**). Interestingly, BPS presented the lowest solvation binding energy value; thus, BPS should bind with lower affinity than BPF and BPA to this configuration, as experimentally demonstrated for the nuclear-initiated pathway (Molina-Molina et al., 2013). These results contrast with the order in biological activity described herein, where we find regulation of ion channel gene expression with at least 100-fold lower concentrations of BPS than BPF. Therefore, a different mechanism to the classic nuclear-initiated event involved in the regulation of ion channel activity and gene expression in β-cells might be implicated in BPS and BPF effects. We then sought to study differences and similarities among E2, BPA, BPS, BPF, and PaPE-1, using what we named rERβ-ΔH12-LBD (PDB code: 1HJ1; (Pike et al., 2001)) dimer complex, and performing long-time (1 µs) molecular dynamics simulations **(Figure 5A)**. In the rERβ-ΔH12-LBD complex, binding of the antiestrogen ICI 164,384 abrogates the association between H12 and the remainder of the LBD, and inhibits both of ER’s transactivation functions (AF1 and AF2) (Pike et al., 2001). BPA does not stabilize ERα in a conformation that initiates nuclear events because BPA does not stabilize H12 (Delfosse et al., 2012). Then, simulating binding to rERβ-ΔH12-LBD dimer complex should be convenient to study extranuclear-initiated events.

**Figure 5.**
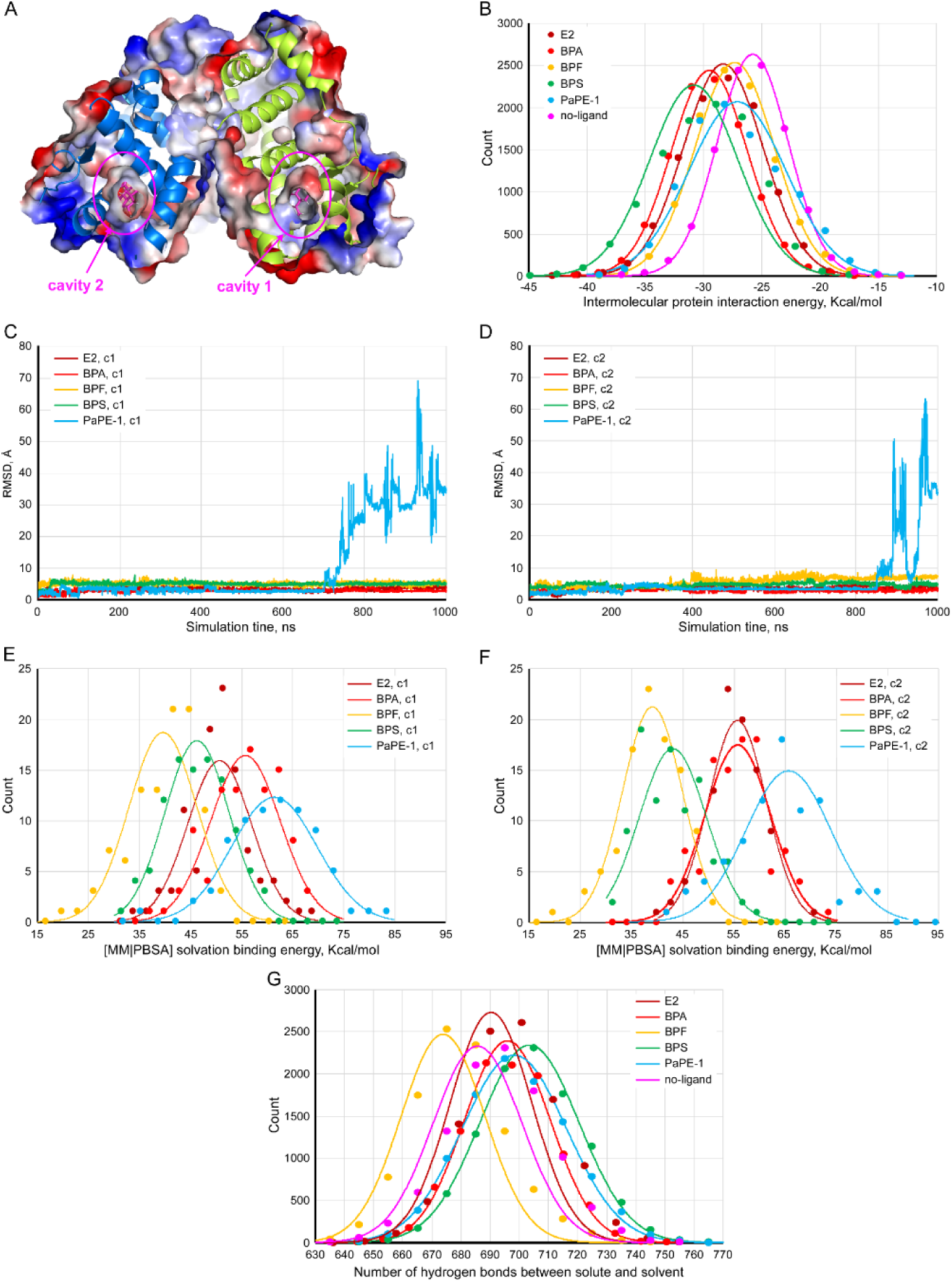
Molecular Dynamics. Analysis of trajectories, MM/PBSA solvation binding energies, and intermolecular interaction energies for the rERβ-ΔH12-LBD dimer from the data generated by MD simulations for 1 µs. **(A)** rERβ-ΔH12-LBD dimer secondary structure and electrostatic surface. The LBD cavity has been cut to show a bound ligand inside the structure. In the following panels, *c1* refers to cavity *1* and *c2* refers to *cavity 2*. **(B)** Frequency distributions of the intermolecular protein interaction energy for the subunits of the rERβ-ΔH12-LBD dimer in the presence of different ligands in each LBD cavity. **(C and D)** trajectories of the ligands (RMSD, Å) initially docked in cavity 1 **(C)** and 2 **(D)** of the LBD. **(E and F)** Frequency distributions of the MM/PBSA solvation binding energy values of each ligand attached to cavity 1 **(E)** and 2 **(F). (G)** Frequency distribution of the number of H-bonds between the solute (protein) and the solvent. A Gaussian curve overlaps discrete data. The legends included within each panel indicate the different ligands analyzed.

Dimerization has been demonstrated to be necessary for ERs-mediated extranuclear responses (Razandi et al., 2004; Levin and Hammes, 2016). In our model, BPA, BPS and E2 showed similar frequency distributions of intermolecular protein interaction for both subunits of the dimers, whereas BPF and PaPE-1 presented lower frequency distribution (**Figure 5B**). This suggests a higher stability of the dimers with BPA, BPS and E2. Trajectories of the ligands docked in the LBD cavities (RMSD, Å) are similar except for PaPE-1 (**Figure 5C**), which left the open cavity of the LBD in both subunits after 700 ns (**Figure 5C**) and 840 ns (**Figure 5D**). For E2 and bisphenols, rearrangements rarely exceed 6 Å, indicating that the movement of the ligands into the cavity is limited, even when H12 is not closing the cavity. Notably, PaPE-1 MM/PBSA solvation binding free energy values, 60 kcal/mol (**Figure 5E**) and 65 kcal/mol (**Figure 5F**), indicate that this compound binds to the protein strongly than bisphenols and E2. On the other hand, bisphenols and E2 remain inside the cavity throughout the entire 1 µs molecular dynamics simulation. We found very similar MM/PBSA solvation binding free energy values for E2 and BPA (around 55 kcal/mol), whereas BPS and BPF presented lower values (45 and 38 kcal/mol, respectively) (**Figure 5E,F**). The solvation binding energies correlated with the number of hydrogen bonds between the protein and the solvent (**Figure 5G)**. BPF presented the lowest solvation binding energy (**Figure 5E,F**) as well as the lowest number of hydrogen bonds (around 680 H-bonds, **Figure 5G**). These results indicate that BPF and PaPE-1 stabilize the ERβ-LBD dimer to a lesser extent than E2, BPA and BPS, which correlates with the lower biological activity observed for BPF and PaPE-1 in the present study. Therefore, we hypothesize that binding of bisphenols to the ERs induces a conformational change that favors an extranuclear-initiated action after dimerization of ERβ.

## DISCUSSION

In the present study we found that acute and long-term exposure of primary male mouse β-cells to BPS and BPF led to alterations in β-cell physiology across different levels of biological complexity, including K_ATP_ channel activity, Na^+^, K^+^ and Ca^2+^ channel subunits expression, and glucose-stimulated insulin release.

Although BPS and BPF have been used as alternatives to BPA, these chemicals may share some of the effects induced by BPA due to their similar structure (Malaise et al., 2020; Mustieles et al., 2020) or induce different effects to BPA (Kolla et al., 2018). We observed that, similarly to BPA, BPS and BPF enhanced GSIS either after acute treatment (1 h) or upon exposure for 48 h.

The acute effect occurs at the two doses tested, 1 nM and 1 µM, and showed glucose dependence. Bisphenols had no effects at a non-stimulatory glucose concentration (3 mM), had a moderate effect at intermediate glucose concentration (8.3 mM), and had a stronger effect at high glucose (16.7 mM). This potentiation was likely a consequence of the blockade of K_ATP_ channels demonstrated using patch-clamp experiments in the cell-attached mode. In β-cells, K_ATP_ channels control the resting membrane potential. As a result of glucose metabolism, the rise in the ratio ATP/ADP blocks K_ATP_ channels and depolarizes the plasma membrane, thus initiating the electrical activity in burst of action potentials that culminates in insulin release. Here we show that BPS- and BPF-induced blockade of K_ATP_ channels seems to potentiate the effect of glucose on insulin secretion, which leads to insulin hypersecretion.

The rapid BPA-induced potentiation of GSIS has been demonstrated in primary mouse and human β-cells (Soriano et al., 2012). In addition, oral BPA administration rapidly altered insulin and C-peptide levels in blood of adult individuals (Stahlhut et al., 2018; Hagobian et al., 2019). As in the present work BPS and BPF act akin to BPA in β-cells, an analogous effect on human cells might be expected. It is difficult to predict how this rapid action relates to the development of metabolic disorders. Pancreatic β-cells acutely exposed to bisphenols secrete more insulin than untreated cells, which may result in supraphysiological insulin signaling in some target tissues, such as adipose tissue.

Although human evidence are still scarce, BPS has been linked to T2D (Ranciere et al., 2019), while BPS and BPF urine levels have been associated with the prevalence of obesity in children (Jacobson et al., 2019; Liu et al., 2019). In any case, bisphenol-induced insulin hypersecretion may be one of the altered processes contributing to insulin resistance, which represents a risk factor for both T2D and obesity (Alonso-Magdalena et al., 2006; Corkey, 2012; Erion and Corkey, 2017). BPS and BPF trigger their rapid actions at concentrations as low as 1 nM. In the presence of 16.7 mM glucose, both 1 nM BPS and BPF increased insulin secretion. However, in the presence of 8.3 mM glucose 1 nM BPS potentiated GSIS, while 1 nM BPF did not affect insulin release. These findings indicate that BPF has a slightly lower potency than BPS, which is manifested by the lack of BPF effect at 1 nM on K_ATP_ channel activity. The difference in potency seems to be small since 10 nM BPF blocked K_ATP_ channel activity to a similar extent as 1 nM BPS. Remarkably, bisphenol-induced blockade of K_ATP_ channels was abolished in cells from BERKO mice. BERKO mice β-cells do not express ERβ (Boronat-Belda et al., 2020). Our previous studies demonstrated that 1 nM of E2, DPN, or BPA similarly affected K_ATP_ channel activity (Soriano et al., 2009; Soriano et al., 2012). These data suggest that ERβ activation blocks K_ATP_ channels and that binding to ERβ may mediate the acute action of bisphenols. It is unlikely that this fast response, reached in only 10 minutes, depends on transcriptional regulation; on the contrary, it most likely relies on extranuclear-initiated pathways involving ERβ. Our findings with PaPE-1 and molecular dynamics as well as the existence of a pool of ERβ outside the nucleus of mouse β-cells (Alonso-Magdalena et al., 2008) support this statement. Designed to selectively trigger extranuclear-initiated pathways, PaPE-1 was obtained after chemical rearrangement of key elements of the original steroid structure of E2 so that its ER binding affinity was considerably reduced (Madak-Erdogan et al., 2016). These modifications were performed by substituting the B-ring of the steroid and methylating the positions 2 and 6 of the A-ring, which prevents the formation of key hydrogen bonds within the ligand binding domain (Madak-Erdogan et al., 2016). Similar methylations are observed in tetramethyl BPF (TMBPF), which had no estrogenic effect as assayed by E-SCREEN and it has been proposed as a safer substitute of BPA (Soto et al., 2017). Here, PaPE-1 blocked K_ATP_ channels in islet cells from WT but not from BERKO mice, which indicates that PaPE-1 and bisphenols activate a similar pathway.

How can bisphenols trigger a rapid effect at low nanomolar concentrations when their affinity for ERβ is within the micromolar range? It is important to bear in mind that the maximum response to a ligand does not depend exclusively on the receptor affinity. The efficacy of the conformational change needed to initiate the signaling cascade as well as the coupling to other signaling proteins also play key roles in the ligand-receptor response (Colquhoun, 1998). Even though the details of the whole pathway from ERβ activation to K_ATP_ closure is not completely known, it has been shown that 1 nM E2 closes K_ATP_ channels through an extranuclear-initiated pathway that involved ERβ, membrane guanylate cyclase, cGMP formation and protein kinase G activation (Ropero et al., 1999; Soriano et al., 2009). The efficacy of this pathway is extremely high as explained below. In addition to the control of the β-cell resting membrane potential, K_ATP_ channels determine the electrical resistance of the β cell membrane (Ashcroft, 2005). When K_ATP_ channels are open, the electrical resistance is low, whereas the resistance is high when these channels are closed. The membrane potential follows Ohm’s law, being the product of the electrical resistance of the membrane by the current running across it. This means that, when extracellular glucose is high, K_ATP_ channels are mostly closed and membrane resistance is high. Hence, a small change in current will elicit membrane depolarization, potentiation of electrical activity, and insulin secretion (Ashcroft, 2005). Our results suggest that bisphenol-induced K_ATP_ channel blockade may lead to enough change in current that will culminate with increased insulin secretion at high glucose. A similar phenomenon is observed with the incretin GLP-1, which acts as an effective secretagogue only when glucose concentrations are stimulatory and a high percentage of K_ATP_ channels are already closed (Holz et al., 1993). Therefore, low doses of bisphenols will be mainly effective under conditions of decreased K_ATP_ channel activity, as seen in the postprandial state. Accordingly, we show that bisphenols are effective insulin secretagogues only when glucose levels are high. Besides their acute effects, longer treatment with bisphenols elicited changes in gene expression and GSIS. As already mentioned, insulin release is a consequence of the electrical activity of pancreatic β-cells, which is determined by the expression of ion channels as well as their biophysical characteristics. Both BPS and BPF decreased the expression of *Cacna1e, Kcnma1* and *Scn9a*, which encode essential subunits of Ca_v_2.3, K_Ca_1.1, and Na_v_1.9 channels. BPS decreased the expression of all channel subunits analyzed at 1 nM, while its effect was lower at 100 nM and 1 µM, which suggests an NMDR relationship. BPF, however, needed higher doses (at least 100 nM) to change channel subunits expression. Therefore, BPS effects on gene expression were 100- and 1000-fold stronger than BPF.

Changes in ion channel expression by 1nM BPA during 48 hours enhanced GSIS (Villar-Pazos et al., 2017; Martinez-Pinna et al., 2019). Here, BPS treatment for 48 h increased GSIS at 1 nM and 1 µM in the presence of 8.3 mM glucose, but only 1 nM BPS was effective in the presence of 16.7 mM glucose. This was surprising and it may indicate the existence of a BPS-triggered mechanism that depends on glucose concentration. A similar effect was described for BPA (Villar-Pazos et al., 2017), in which BPA exposure for 48 h decreased exocytosis at low glucose (5.6 mM) but increased exocytosis at high glucose concentrations (11 mM). While these findings suggest the existence of a crosstalk between BPA and glucose signaling effects on the exocytotic machinery, the existence of such crosstalk is yet to be elucidated. Exposure to BPF enhanced GSIS only at 1 µM, the same concentration at which gene expression occurred. These results emphasize the different potencies observed between BPS and BPF.

Our results in BERKO mice indicate that both BPS and BPF effects on gene expression are mediated by ERβ. BPF acts within the micromolar range, which is compatible with its ERβ affinity (see below). On the other hand, our data also suggest that 1 nM BPS acts through ERβ, which is surprising if we consider that BPS binds to ERβ and activates the classic nuclear-initiated pathway at higher concentrations. In vitro bioassays using the stably transfected HELN-hERβ cell line, which contains a luciferase gene driven by an ERE under the control of hERβ, have clearly demonstrated that BPA, BPS and BPF behaved as full hERβ agonists with potencies in the following order: BPA>BPF>BPS (Molina-Molina et al., 2013). Additionally, whole-cell competitive binding assays using the same cell line showed IC_50_ values of 0.21±0.01 nM (E2), 401±126 nM (BPA), 1452±261 nM (BPF), and 3452±878 nM (BPS) (Molina-Molina et al., 2013). Although our findings with BPF are compatible with this classic model, this does not seem to be the case for BPA and BPS.

We pointed out in the first part of the Discussion that low doses of bisphenols can signal through extranuclear-initiated pathways in β-cells. We showed that, like bisphenols, PaPE-1 is an agonist that uses this extranuclear pathway to decrease K_ATP_ channel activity in an ERβ-dependent manner. Then, it is possible that an extranuclear-initiated pathway may be implied in bisphenols action to explain their effects at nanomolar concentrations.

As already discussed, the efficacy of bisphenol response would depend on the interaction between ERβ and other proteins involved in extranuclear signaling. Molecular dynamics indicated that dimerization may be important and may explain, at least in part, why BPA and BPS are more potent than BPF. Dimerization is a requisite for ER extranuclear signaling (Razandi et al., 2004; Levin and Hammes, 2016) and its role deserves further research in the case of bisphenols and other xenoestrogens. Extranuclear signaling by nuclear receptors is a complex phenomenon and there are very few data showing the activation of such extranuclear pathways by endocrine-disrupting chemicals, including bisphenols (Marino et al., 2012; Vinas and Watson, 2013; Nadal et al., 2018). ERs acting through this pathway do not directly engage DNA to regulate transcription but induce non-nuclear signaling cascades that may lead to transcriptional regulation. Extranuclear ERs interact with a plethora of signaling proteins associated to the plasma membrane or present in the cytosol, such as G proteins and other receptors and kinases involved in extranuclear-initiated signaling triggered by estrogens (Levin and Hammes, 2016). These interactions may amplify bisphenol response via extranuclear ERβ as it has been shown for adrenergic and cholinergic receptors, which respond to ultralow concentrations of ligand within the femtomolar range (Civciristov et al., 2018). Thus, it is necessary to further study bisphenol-activated extranuclear-initiated ER signaling pathways to better understand the efficacy of the response. This information is urgently needed to develop improved testing methods for extranuclear-initiated effects as well as to fully explain how low doses of estrogenic endocrine-disrupting chemicals affect several biological processes.

## CONCLUSIONS

Both short- and long-term exposure to BPS and BPF increases glucose-induced insulin release, which is a risk factor for T2D. A rapid response may be due to the closure of K_ATP_ channels, while a long-term response seems to be via regulation of ion channel gene expression. As K_ATP_ channel activity, gene expression of ion channel and insulin release are endpoints relatively easy to be measured, we propose they should be considered key events to assess the potential hazards of bisphenols.

In line with previous work with ERβ agonists and BPA, our findings with BERKO mice and PaPE-1 suggest that bisphenols act as ERβ agonists and activate an extranuclear-initiated pathway. ERβ affinity for BPA and BPS cannot easily explain the biological effects described in the present work and in previous reports. Our data with acute exposure indicate that efficacy may be more important than affinity to explain effects at low doses. More experimental data on dimerization and interaction of ERβ with other signaling molecules in its vicinity are needed to fully understand effects at low doses of bisphenols, especially on gene transcription. In any case, our data support that these bisphenols are not a safe alternative to BPA.

## AUTHOR INFORMATION

### Author Contributions

**LM**: Conceptualization, Supervision, Investigation, Formal Analysis, Visualization, Writing - Review & Editing **JMP**: Conceptualization, Investigation, Formal Analysis, Visualization, Writing - Review & Editing **MCM**: Investigation, Formal Analysis, Visualization, Writing - Review & Editing **RSdS**: Investigation, Formal Analysis, Writing - Review & Editing **RMMG**: Investigation, Formal Analysis, Writing - Review & Editing **SS**: Investigation, Formal Analysis, Visualization, Writing - Review & Editing **IQ**: Visualization, Resources, Funding Acquisition, Writing - Review & Editing **J-AG**: Resources, Writing - Review & Editing **JAE**: Conceptualization, Investigation, Visualization, Resources, Funding Acquisition, Writing - Review & Editing **AN**: Conceptualization, Supervision, Visualization, Resources, Funding Acquisition, Writing - Original Draft, Project Administration. All authors have given approval to the final version of the manuscript.

## ACKNOWLEDGMENT

The authors thank Maria Luisa Navarro, Salomé Ramon, and Beatriz Bonmati Botella for their excellent technical assistance. We are grateful to the Cluster of Scientific Computing (http://ccc.umh.es/) of the Universitas Miguel Hernández for providing computing facilities.

