## Supplementary Data for "Bisphenol-S and Bisphenol-F alter mouse pancreatic β-cell ion channel expression and activity and insulin release through an estrogen receptor ERβ mediated pathway"

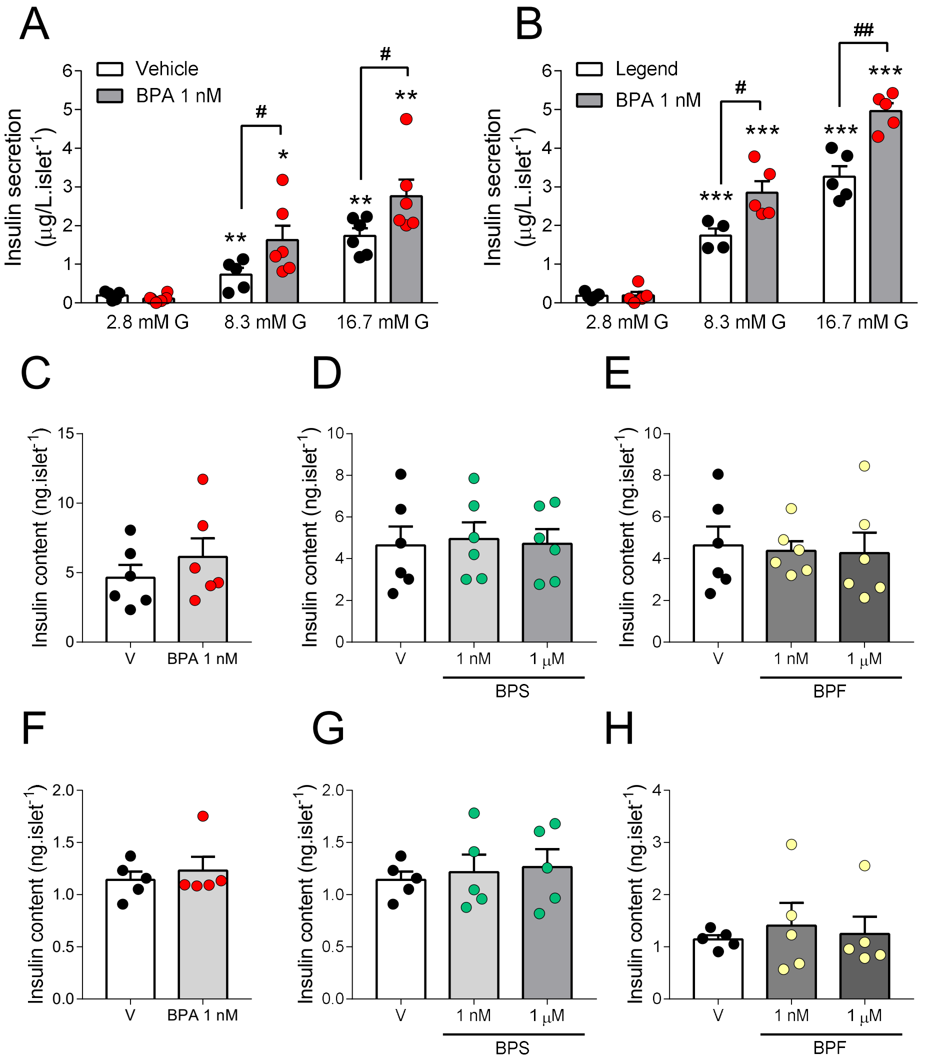


**Figure S1. Insulin secretion and content upon treatment with bisphenols**. **(A and B)** Insulin secretion was measured at 2.8, 8.3 and 16.7 mM glucose in islets from C57BL/6J mice treated *ex vivo* with vehicle (control; black circles and white bars) or 1 nM BPA (red circles and light grey bars). **(C-H)** Insulin content was measured after GSIS of the experiments described in the **Figure 1**. Mouse islets from C57BL/6J mice treated *ex vivo* with vehicle (control; black circles and white bars), 1 nM BPA (**A and F**; red circles), BPS (**D and G**; green circles) or BPF (**E and H**; yellow circles). Insulin content was measured by ELISA. Data are shown as means ± SEM of six independent islet preparations isolated on three different days: *$p \leq 0.05$, **$p \leq 0.01$, ***$p \leq 0.001$ vs 2.8 mM; $^{\#}p \leq 0.05$, $^{\#\#}p \leq 0.01$, $^{\#\#\#}p \leq 0.001$ comparisons indicated by bars (one-way ANOVA); $^{\&}p \leq 0.05$ (Student's t-test).

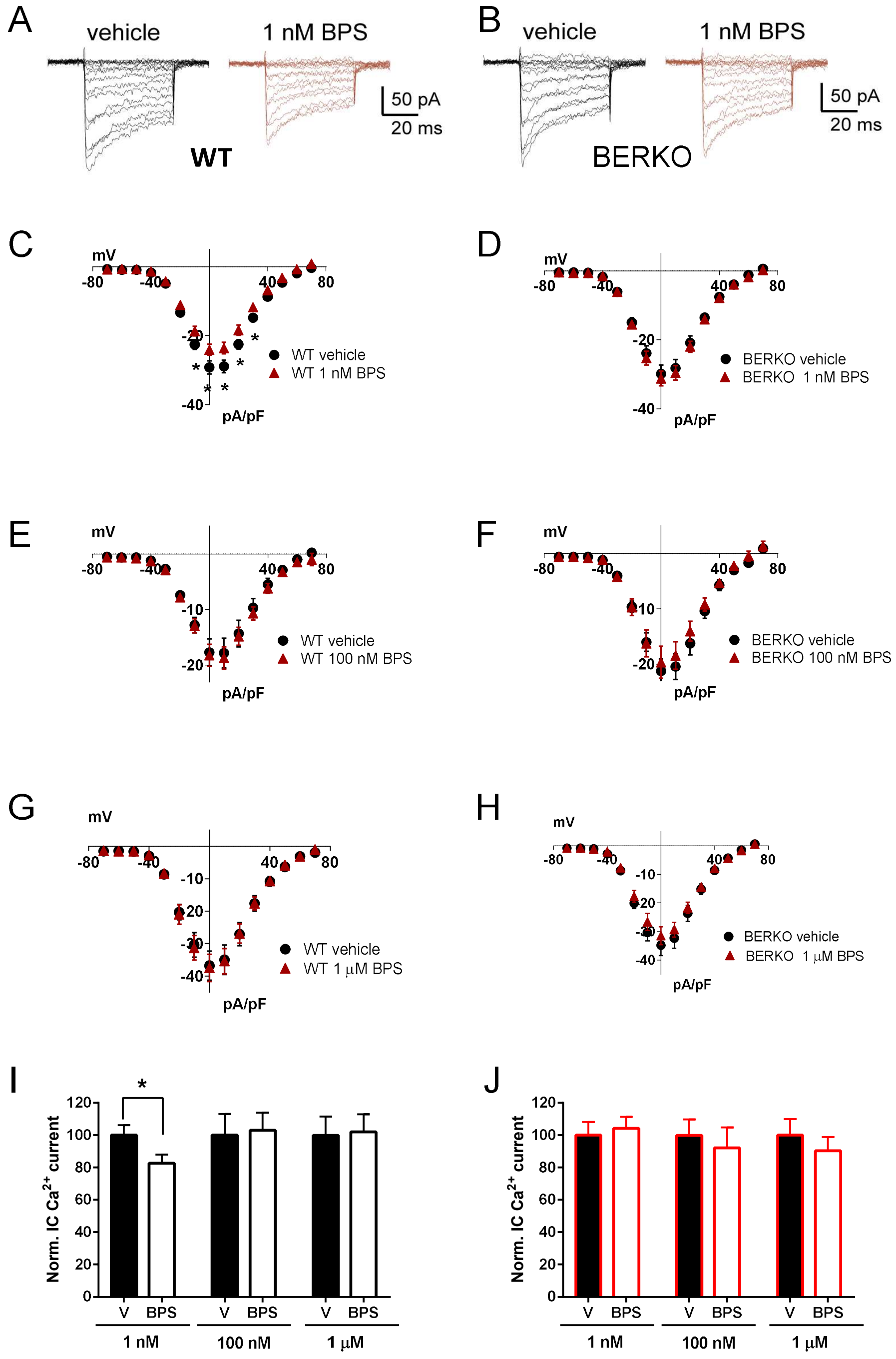


**Figure S2. Low doses of BPS reduce whole-cell $Ca^{2+}$ currents via ERβ in *β-cells*.** (**A and B)** Representative recordings of whole-cell $Ca^{2+}$ currents in response to depolarizing voltage pulses (-60 to +70 mv from a holding potential of -70 mV, 50 ms duration) in isolated β-cells from wild type (WT, **A**) or BERKO (**B**) mice upon treatment with vehicle (left black traces) and 1 nM BPS (right red traces). **(C-H)** Average relationship between $Ca^{2+}$ current density ($Ca^{2+}$ currents in pA normalized to the cell capacitance in pF) and the voltage of the pulses in wild-type (WT, **C**, **E and G**) and BERKO (**D**, **F and H**) control cells (black circles) and cells treated (red triangles) with 1 nM BPS (**C** and **D**), 100 nM BPS (**E and F**) or 1 µM BPS (**G and H**). **(I and J)** Average normalized values of current density evoked at 0 mV obtained from the I–V relationship shown in WT mice **(C, E and G)** and the BERKO littermates **(D, F and H)**. The effect of BPS was measured after 48 h of incubation. Data are shown as means ± SEM of the number of cells recorded in WT (n=10-15 cells) and BERKO (n=7-15 cells) mice. These cells were isolated from six mice on at least three different days: *$p \leq 0.05$ vs control (one-way ANOVA).

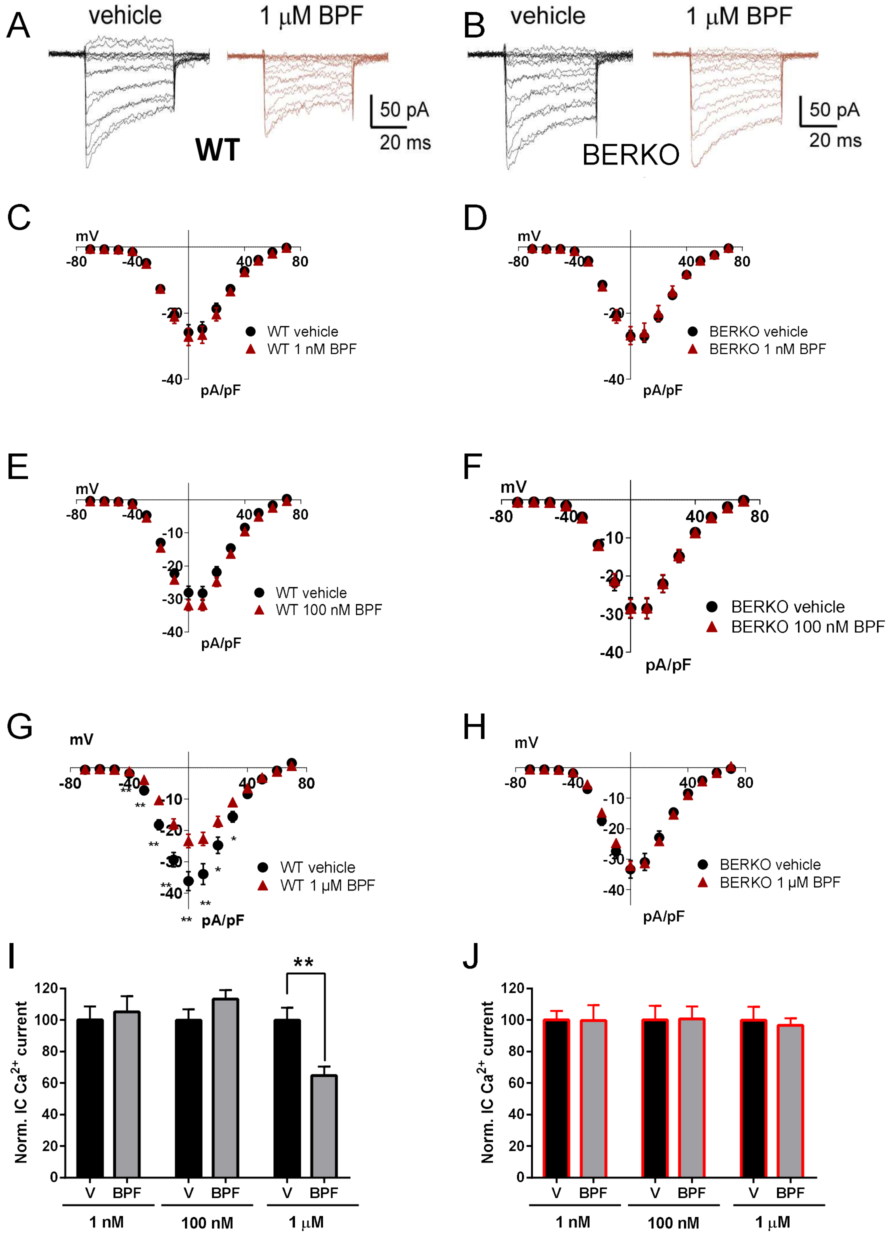


**Figure S3. High doses of BPF reduce whole-cell $Ca^{2+}$ currents via ERβ in *β-cells*.** **(A and B)** Representative recordings of whole-cell $Ca^{2+}$ currents in response to depolarizing voltage pulses (-60 to +70 mv from a holding potential of -70 mV, 50 ms duration) in isolated β-cells from wild type (WT, **A**) or BERKO (**B**) mice upon treatment with vehicle (left black traces) and 1 μM BPF (right red traces). **(C-H)** Average relationship between $Ca^{2+}$ current density ($Ca^{2+}$ currents in pA normalized to the cell capacitance in pF) and the voltage of the pulses in wild-type (WT, **C**, **E and G**) and BERKO (**D**, **F and H**) control cells (black circles) and cells treated (red triangles) with 1 nM BPF (**C and D**), 100 nM BPF (**E and F**) or 1 μM BPF (**G and H**). **(I and J)** Average normalized values of current density evoked at 0 mV obtained from the I–V relationship shown in WT mice **(C, E and G)** and the BERKO littermates **(D, F and H)**. The effect of BPF was measured after 48 h of incubation. The methodology for patch-clamp recordings of voltage-gated $Ca^{2+}$ currents is the same as the one described in Figure S2. Data are shown as means ± SEM of the number of cells recorded in WT (n=13-21 cells) and BERKO (n=9-23cells) mice. These cells were isolated from six mice on at least three different days: *$p \leq 0.05$ vs control (one-way ANOVA).

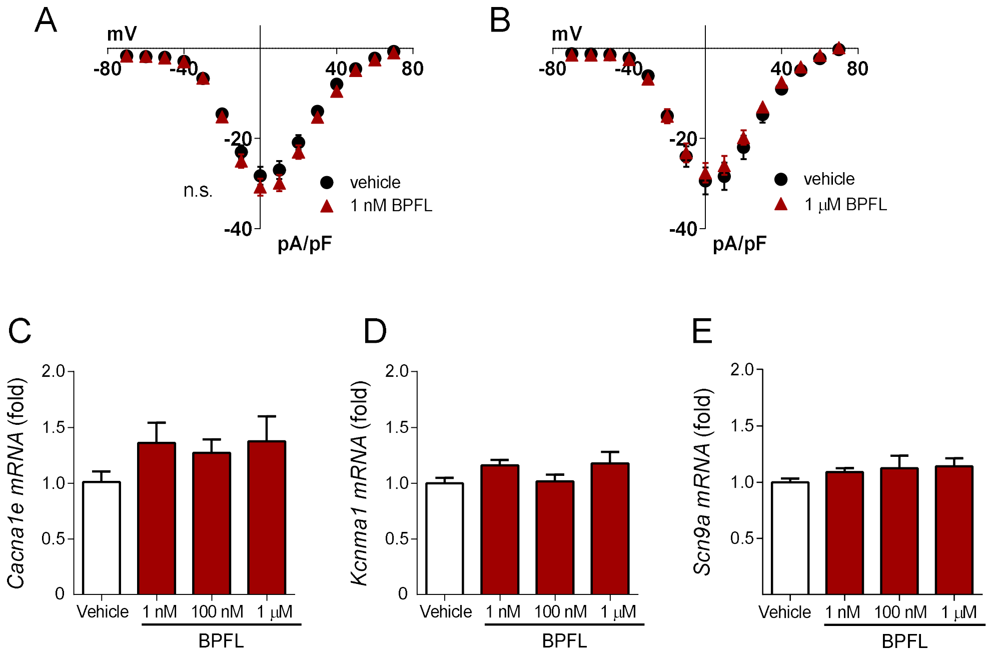


**Figure S4. Bisphenol FL does not change whole-cell $Ca^{2+}$ currents or *Cacna1e*, *Kcnma1* and *Scn9a* mRNA expression in mouse islets.** (**A and B**) Average relationship between $Ca^{2+}$ current density ($Ca^{2+}$ currents in pA normalized to the cell capacitance in pF) and the voltage of the pulses (-60 to +70 mv from a holding potential of -70 mV, 50 ms duration) in isolated β-cells treated *in vitro* with vehicle (black circles) or with 1 nM BPFL (**A**; red triangles) or 1 µM BPFL (**B;** red triangles). The effect of BPFL was measured after 48 h of incubation. Data are shown as means ± SEM of the number of cells recorded in vehicle (n=8-10 cells) and BPFL (n=7-9 cells). These cells were isolated from six mice on at least three different days (**C-E**) mRNA expression of *Cacna1e* **(C)**, *Kcnma1* **(D)** and *Scn9a* **(E)** was measured in islets from C57BL/6J mice treated *ex vivo* with vehicle (white bars) or BPFL (red bars) at 1 nM, 100 nM, and 1 µM for 48 h. mRNA expression was measured by qRT-PCR and normalized to the housekeeping gene *Hprt1,* and is shown as fold vs. mean of the controls. Data are shown as means ± SEM of: four to twenty independent samples from up to twenty islets preparations isolated on at least three different days (one-way ANOVA).

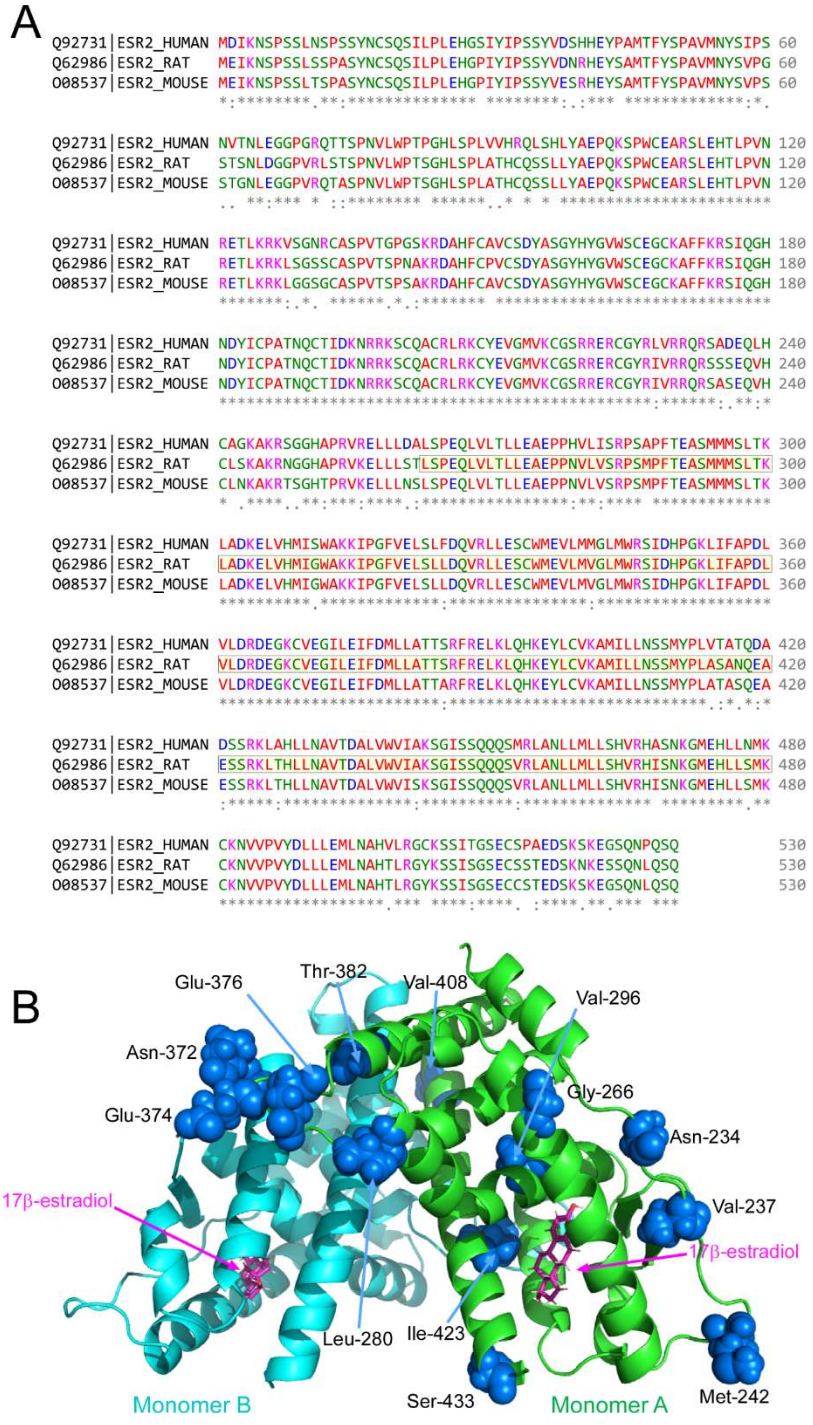


**Figure S5. Sequence alignment of human, rat and mouse LBD-ERβ.** (**A**) Multiple sequence alignment of human, rat and mouse ERβ. A yellow box indicates the region of the protein corresponding to the LBD and whose structure has been resolved from x-ray diffraction data. An orange box locates the human Cys-339 that can be palmitoylated. (**B**) Secondary structure of the rat ERβ-LBD dimer (PDB ID 1HJ1) that includes a 17β-estradiol molecule in the ligand-binding cavity of each subunit. Monomer A (green) highlights the different amino acids (spheres, Asn-234, Val-237, Met-242, Gly-266, Leu-280, Val-296, Ala-369, Ser-370, Asn-372, Glu-374, Glu-376, Thr-382, Val-408, Ile-423, Ser-433) in the human isoform.

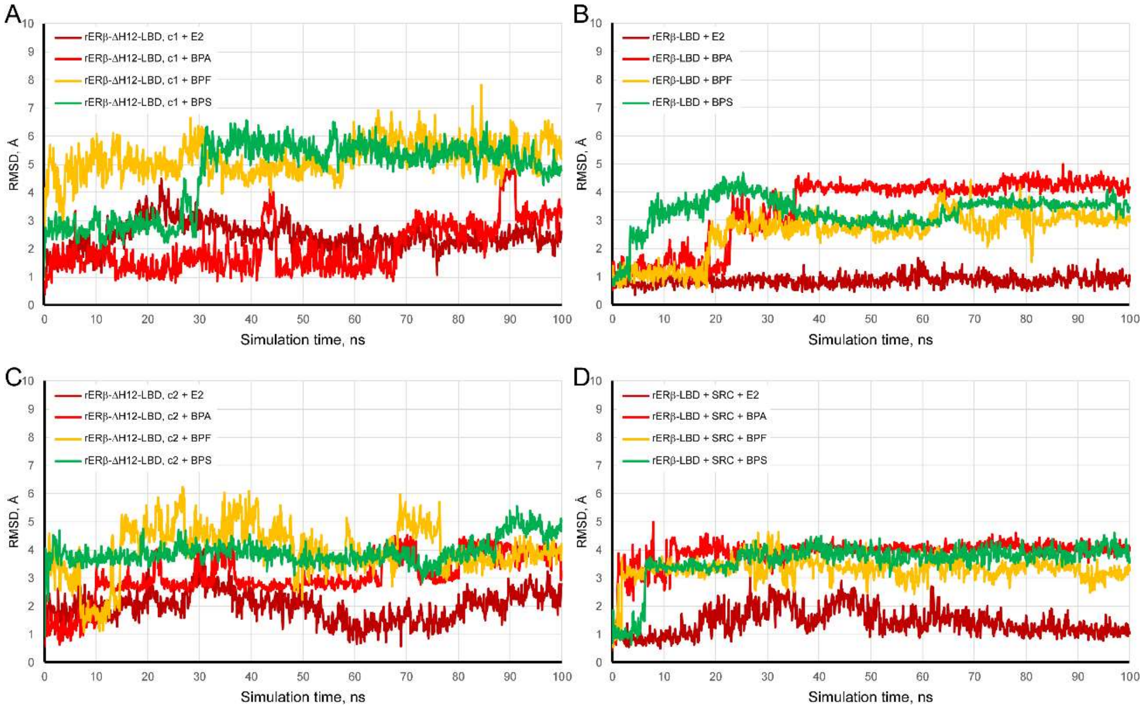


**Figure S6. Analysis of trajectories for molecular dynamics simulations of LBD-ER in complex with different ligands. A and C** Trajectories (RMSD, Å) of the different ligands initially docked to the main cavity of the open rERβ-ΔH12-LBD dimer. Cavity 1 (c1, **A**) and cavity 2 (c2, **C**). **B** and **D** Trajectories (RMSD, Å) of the different ligands docked to the closed cavity of the rERβ-LBD monomer in the absence **(B)** or presence **(D)** of the SRC co-activating peptide, respectively. The legends included within each panel indicate the different ligands analyzed.

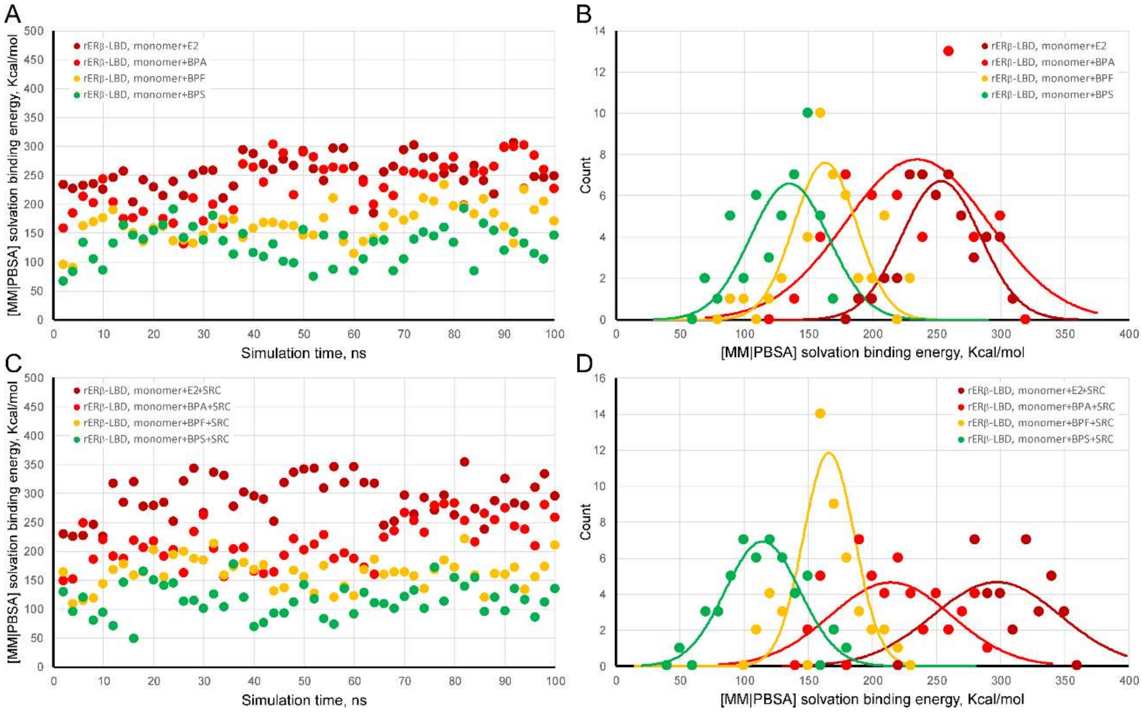


**Figure S7. Calculated MM/PBSA solvation binding energy for bound E2 and bisphenols to the H12 closed rERβ-LBD alone or in the presence of the Src peptide. (A and C)** Calculated MM/PBSA solvation binding energy values of each ligand attached to LBD cavity alone **(A)** or in the presence of the Src peptide **(C)**. **(B and D)** Frequency distributions of the values shown in **(A)** and **(C)**, respectively. A Gaussian curve overlaps discrete data. The legends included within each panel indicate the different ligands analyzed.
